# The Prorenin Receptor and its Soluble Form Contribute to Lipid Homeostasis

**DOI:** 10.1101/2020.01.27.922138

**Authors:** Eva Gatineau, Gertrude Arthur, Audrey Poupeau, Kellea Nichols, Brett T. Spear, Gregory Graf, Ryan Temel, Frédérique Yiannikouris

**Affiliations:** Department of Pharmacology and Nutritional Sciences, University of Kentucky, Lexington, KY 40536; Department of Microbiology, Immunology and Molecular Genetics, University of Kentucky, Lexington, KY 40536; Department of Pharmaceutical Sciences, University of Kentucky, Lexington, KY 40536; Cardiovascular Research Center and Department of Physiology, University of Kentucky, Lexington, KY 40536

**Keywords:** PRR, sPRR, liver, adipose tissue, lipid homeostasis

## Abstract

Obesity is associated with alterations in hepatic lipid metabolism. We previously identified the prorenin receptor (PRR) as a potential contributor to liver steatosis. Therefore, we aimed to determine the relative contribution of PRR and its soluble form, sPRR, to lipid homeostasis. PRR-floxed male mice were treated with an adeno-associated virus with thyroxine-binding globulin promoter driven Cre to delete specifically PRR in hepatocytes (Liver PRR KO mice). Hepatic PRR deletion did not change the body weight but increased liver weights. Liver PRR KO mice exhibited higher plasma cholesterol levels and lower hepatic LDLR protein than control mice. Surprisingly, hepatic PRR deletion elevated hepatic cholesterol, and up-regulated hepatic SREBP2 and HMG CoA-R genes. In addition, hepatic PRR deletion increased plasma sPRR levels. *In vitro* studies in Hep-G2 cells demonstrated that sPRR treatment up-regulated SREBP2 suggesting that elevated plasma sPRR could contribute to hepatic cholesterol biosynthesis. Interestingly, PPARγ, PRR and total sPRR were elevated in the adipose tissue of Liver PRR KO mice suggesting that elevated plasma sPRR originated from the adipose tissue. In 3T3-L1 cells, sPRR treatment up-regulated PPARγ indicating that sPRR stimulates master regulator of adipocyte differentiation. Overall, this work support a new role for sPRR in lipid metabolism and adipose tissue – liver crosstalk.

## Introduction

The obesity epidemic, currently affecting more than 35% of the population in the United States, is associated with several deleterious changes in lipid metabolism, including increased serum lipids, and glucose/insulin homeostasis. Hyperlipidemia is a risk factor for cardiovascular disease (CVD) and is estimated to be responsible for more than half of cardiovascular mortality (1–3).

The prorenin receptor (PRR) is a component of the renin angiotensin system involved in the regulation of blood pressure and fluid volume. PRR exists in three different forms: a full-length integral transmembrane protein, a soluble PRR (sPRR) and a truncated form composed of a transmembrane and a cytoplasmic domain (4–7). Increasing evidence suggests that the function of PRR is not restricted to blood pressure control (8–10). PRR can also interact with the membrane receptor complex of the Wnt/β-catenin pathway or with the vacuolar-type H-ATPase, two crucial pathways involved in tissue and organ development (8, 11–14). Evidence also indicates that both PRR and sPRR play role in obesity, energy and lipid metabolism (15–18). Indeed, we previously showed that adipose PRR KO prevented the development of obesity and drastically decreased fat mass (15). Paradoxically, the deletion of PRR in adipose tissue increased plasma sPRR and induced lipid redistribution in the liver that led to liver steatosis (15). Moreover, sPRR and PRR was elevated in the liver of adipose PRR KO mice (19) suggesting that the elevation of plasma sPRR likely originated from the liver.

Lu *et al.*, silenced PRR in hepatocytes and found that PRR deletion decreased cellular LDL uptake and reduced SORT1 and LDLR proteins levels (16). In mice, the knockdown of PRR in liver, using antisense oligonucleotides (ASO), impaired hepatic LDL clearance (17). However, plasma sPRR levels were not quantified in this study. Thus, the relative contribution of sPRR to lipid homeostasis was not assessed. Therefore, we first aimed to determine whether the deletion of PRR specifically in hepatocytes influenced plasma sPRR levels in mice fed a standard diet using an adeno-associated viral (AAV)-thyroxine binding globulin (TBG) promoter-Cre recombinase vector administered to PRR “floxed” mice (Liver PRR KO). Since we found that the deletion of hepatic PRR increased plasma sPRR levels, we next investigated the tissue origin of this elevation and the relative contribution of sPRR to lipid metabolism in liver PRR KO mice.

### Experimental protocol

#### Animals

All procedures involving animals were conducted with the approval of the University of Kentucky Institutional Animal Care and Use Committee (University of Kentucky IACUC protocol number: 2013-1109) and in accordance with the National Institutes of Health Guide for the Care and Use of Laboratory Animals.

#### AAV experiment

An adeno-associated virus was generated by the Genetic Technologies CORE of the University of Kentucky. Cre recombinase was driven by a liver-specific promoter, thyroxine-binding globulin (TBG) promoter (*TBG^p^-Cre*). PRR floxed male mice (8 to 12 month old) with loxP sites flanking exon 2 of the PRR gene (*PRR^fl/fl^*) were treated with a single intraperitoneal injection at a dosage of 1.0 × 10^11^ gc virus (Liver PRR KO, n=5) or with saline (CTL, n=6). Mice were provided water and normal chow diet ad libitum (18% protein, Global Diet; Teklad Harlan Madison, WI). Three weeks after injection, tissues were harvested, snap frozen in liquid nitrogen, and stored at −80°C. Blood was collected in tubes (4°C) containing EDTA (0.2 mol/L), centrifuged at 5,000 rpm for 10 min, and plasma was stored at −80°C.

#### Quantification of plasma and tissues components

Plasma concentrations of sPRR were quantified using a soluble (Pro)renin Receptor ELISA kit (Immuno-Biological Laboratories Co, Minneapolis, MN). Plasma concentrations of cholesterol and triglyceride were measured using Wako kit (Wako Chemicals, Richmond, VA). Total sPRR contents of epididymal fat (EF) was assessed using soluble (Pro)renin Receptor ELISA kit (Immuno-Biological Laboratories Co, Minneapolis, MN). The antibody anti-PRR of the sPRR ELISA kit recognized the same epitope for sPRR and full-length PRR and therefore assessed total sPRR content in tissue.

### Lipoprotein cholesterol and apolipoprotein distribution

The cholesterol distribution among lipoprotein classes was determined after separation by gel filtration chromatography based upon the method described previously (20). An aliquot of plasma was diluted in 0.9% NaCl, 0.05% EDTA/NaN3 and centrifuged at 2000 x g for 10 minutes to remove any particulate debris. The supernatant was transferred to a glass insert contained in a GC vial. After loading the vial into an autosampler set at 4°C (G1329A, Agilent Technologies, Santa Clara, CA), 40 μL of sample was injected onto a Superose 6 10/300 or Superose 6 Increase 10/300 (GE Healthcare Life Sciences, Pittsburgh, PA) chromatography column. Under the control of an isocratic pump (G1310A/B, Agilent Technologies, Santa Clara, CA), the sample was separated at a flow rate of 0.4 ml/min with eluent containing 0.9% NaCl, 0.05% EDTA/NaN3. The column effluent was mixed with total cholesterol enzymatic reagent (C7510, Pointe Scientific, Canton, MI) running at a flow rate of 0.125 mL/min and the mixture was passed through a knitted reaction coil (EPOCOD, Aura Industries Inc., San Diego, CA) in a 37°C H2O jacket. The absorbance of the reaction mixture was read at 500 nm using a variable wavelength detector (G1314F, Agilent Technologies, Santa Clara, CA). The signal was subsequently integrated using Agilent OpenLAB Software Suite (Agilent Technologies, Santa Clara, CA). For apolipoprotein distribution, plasma was separated by gel filtration chromatography as described above and then 3 min fractions were collected between 18 and 48 minutes for further analysis by western blot.

#### Quantification of liver lipids

Liver lipid content was determined based upon the method described by Carr *et al.*,(21). A piece of frozen liver was thawed and minced with a razor blade. Following transfer to a tared 16×100mm glass tube, the wet weight of the tissue was measured using an analytical balance. To extract the lipids from the tissue, 3 ml 2:1 chloroform:methanol (CHCl3:MeOH) was added and incubated at 55°C for 2 hours. After centrifuging the tube at 1,500xg for 10 min, the lipid extract was transferred to a new 16×100 mm glass screw top tube. The tube containing the extracted liver was washed with 2 ml 2:1 CHCl3:MeOH and centrifuged as described above. The lipid extract and wash were combined and the solvent was evaporated under nitrogen at 55°C. The dried lipid extract was dissolved in 6 ml of 2:1 CHCl3:MeOH. After the addition of 1.2 ml dilute H2SO4 (0.05%, v/v), the sample was vortexed for 20 seconds and the phases were separated by centrifugation as described above. The upper aqueous phase was removed and an aliquot (typically 1 ml) of the bottom, lipid-containing organic phase was transferred to a new 16×100 mm glass screw top tube using a volumetric glass pipet. After adding 2 ml 1% Trition-X100 dissolved in CHCl3, the organic solvent was evaporated under nitrogen at 55°C. The dried sample was dissolved in 1 ml water and heated at 60°C for 10 min. After vortexing and centrifuging as above, samples dissolved in 2% Triton-X100/water were analyzed for triglycerides using a Wako kit (Wako Chemicals, Richmond, VA). Standards for the triglycerides assay were created using vegetable oil and were dissolved in 2% Triton-X100/water. Organic-solvent resistant, Teflon lined caps were used to seal the tubes throughout the protocol.

#### Tissue RNA extraction and quantitative RT-PCR

RNA was extracted from liver and EF using the SV Total RNA Isolation System (Promega, Madison, WI). Total RNA was quantified with a NanoDrop 2000 spectrophotometer (Wilmington, DE) and cDNA was synthesized using qscript cDNA SuperMix (Quanta Biosciences, Gaithersburg, MD). Real-time quantitative PCR was performed with PerfeCTa SYBR Green FastMix (Quanta BioSciences, Gaithersburg, MD). All the primers sequences are described in Supplemental Table 1.

#### Western blotting

Protein from frozen liver and EF were extracted in ice-cold Tris buffer enriched with Roche cOmplete cocktail inhibitor with a Geno/Grinder® 2010 (SPEX SamplePrep, Metuchen, NJ) and were submitted to SDS-PAGE on precast polyacrylamide gel (Mini-PROTEAN® TGX™, 4-20%, Bio-Rad Laboratories, Hercules, CA). Proteins were transferred on polyvinylidene difluoride membrane, which was blocked in 5% non-fat dried milk in Tris-buffered saline with 0.1 % Tween 20 (TBST). Membranes were then incubated with anti-PRR antibody (#HPA003156, Sigma, St Louis, MO), anti-LDLR (#ab30532 Abcam, Cambridge, MA), anti-PPARγ (#2435, Cell Signaling Technology, Inc., Danvers, MA), anti-Perilipin (#9349, Cell Signaling Technology, Inc., Danvers, MA), anti-FABP4 (#2120, Cell Signaling Technology, Inc., Danvers, MA), anti-CEBPα (#8178, Cell Signaling Technology, Inc., Danvers, MA), or anti-GAPDH (#5174, Cell Signaling Technology, Inc., Danvers, MA) in TBST 5% non-fat dried milk. Following incubation with HRP-conjugated anti-rabbit secondary antibody (Jackson ImmunoResearch Laboratories, West Grove, PA), proteins were imaged using a Syngene PXi imager (Syngene, Frederick, MD). The levels of proteins were quantified using ImageJ software (National Institutes of Health, Bethesda, MD, USA) and normalized to GAPDH levels. For quantification of lipoprotein in FPLC fractions, 10 μl of pooled fractions were separated on a 4-20% SDS-PAGE gel, transferred to PVDF membranes and incubated overnight with anti-ApoA1 (#K23001R, Meridian Life Science, Inc., Memphis, TN) or ApoB48/100 (#K23300R, Meridian Life Science, Inc., Memphis, TN) antibodies. After incubation with HRP-conjugated anti-rabbit secondary antibody (#A6154, Sigma, St Louis, MO) protein were revealed on an X-ray film and quantified as described above.

#### *In vitro* experiments

Hep-G2 cells (ATCC) were cultured in the presence of Dulbecco’s modified Eagle’s medium (DMEM) supplemented with 10% (v/v) fetal bovine serum and 1% (v/v) of a mixture of penicillin and streptomycin. PRR was silenced by mouse Stealth RNA interference designed for Atp6ap2 gene (PRR) using lipofectamineTM 2000 (life technology), and according to the procedure recommended by the manufacturer. 3T3-L1 cells were cultured in the presence of DMEM supplemented with 10% (v/v) fetal bovine serum and 1% (v/v) of a mixture of penicillin and streptomycin and treated with mouse recombinant sPRR-HisTag (residues 18–276, Genscript) or with vehicle for 24 hours.

#### Statistical Analysis

Data are represented as means ± the standard error of the mean (SEM). Statistical differences between groups were analyzed by one-way ANOVA followed by Holm-Sidak post-hoc analysis for multiple comparisons. For ANOVA models, normality was assessed using the Shapiro-Wilk test, and when p<0.05, response variables were log transformed. Correlations between two variables were assessed using Pearson correlation coefficient. Statistical outliers were identified using Grubbs test (GraphPad QuickCalcs).

## Results

### The deletion of PRR in liver induced hepatomegaly and disturbed lipid homeostasis

Male *PRR*^*fl/fl*^ mice treated with AAV-*TBG*^*p*^-*Cre* (Liver PRR KO) exhibited significantly reduced liver PRR mRNA (Figure 1A) and protein levels (Figure 1B) compared to mice treated with vehicle (CTL). The body weight of Liver PRR KO mice were not significantly different from CTL mice (Figure 1C). The deletion of PRR in liver did not significantly change total fat and lean masses (expressed as percent of body weight, Figure 1D). Moreover, the weights of the retro-peritoneal fat (RPF), epididymal fat (EF) and subcutaneous fat (subc) were not significantly different in Liver PRR KO mice compared to CTL mice (Table 1). Interestingly, liver weights increased significantly in Liver PRR KO compared to CTL mice (Table 1). Additionally, the deletion of hepatic PRR increased the levels of plasma total and free cholesterol (Figure 2A and 2B), and LDL cholesterol (Figure 2D and 2E) and did not change plasma triglycerides levels (Figure 2C). Hepatic LDLR protein levels (non glycosylated form, 95 KDa) decreased significantly in Liver PRR KO mice compared to CTL mice (Figure 3A and 3B).

**Figure 1.**
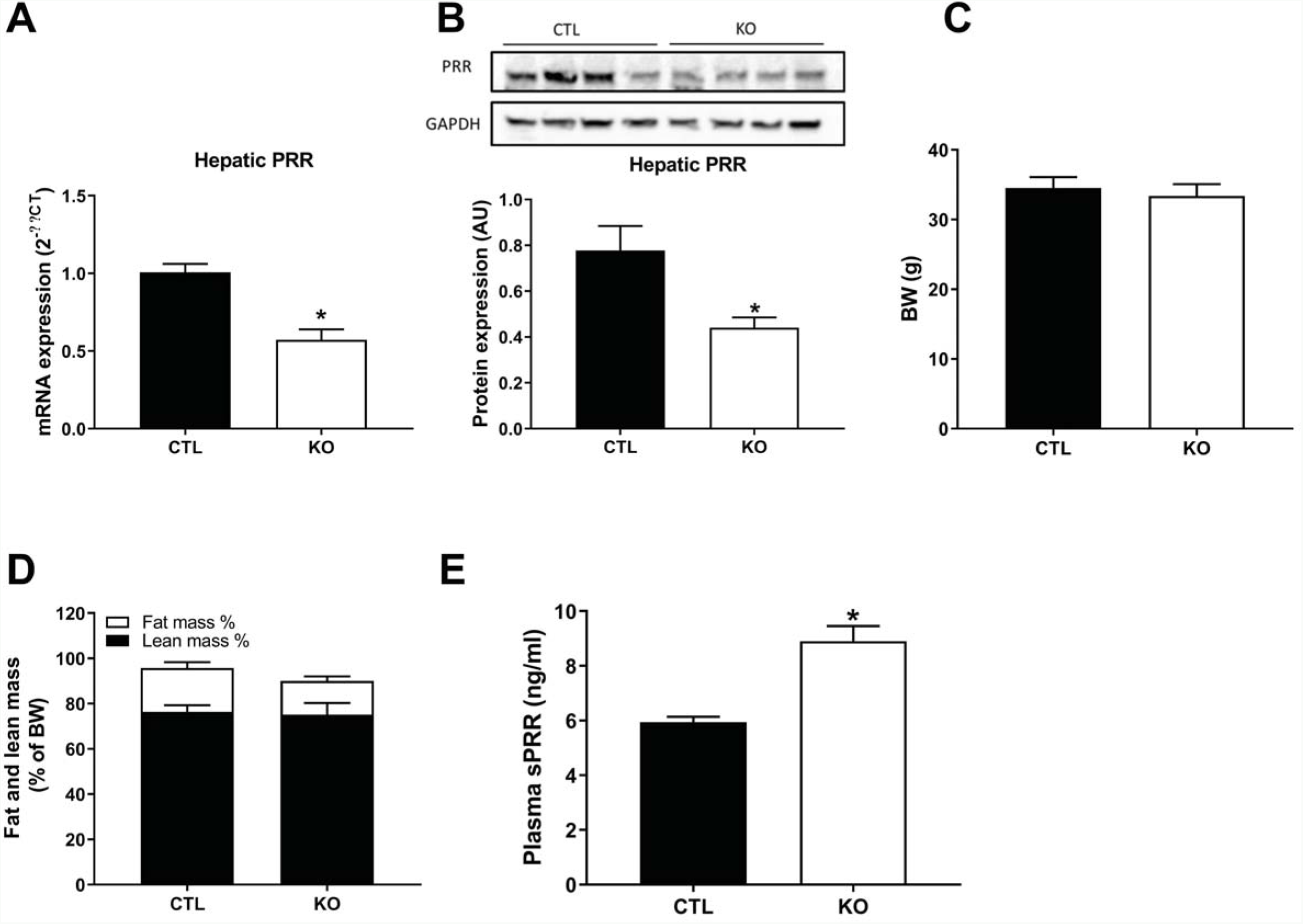
The deletion of hepatic PRR did not change the body weight and the fat mass. (A) Hepatic mRNA expression of PRR in control (CTL) and liver PRR KO male mice (KO). (B) Hepatic protein levels of control (CTL) and liver PRR KO male mice (KO). (C) Body weight of PRR in control (CTL) and liver PRR KO male mice (KO). (D) Body composition of PRR in control (CTL) and liver PRR KO male mice (KO) assessed by EchoMRI. (C: n=6 and KO: n=5). Data are mean±SEM. *P<0.05 compared with vehicle.

**Table 1.**
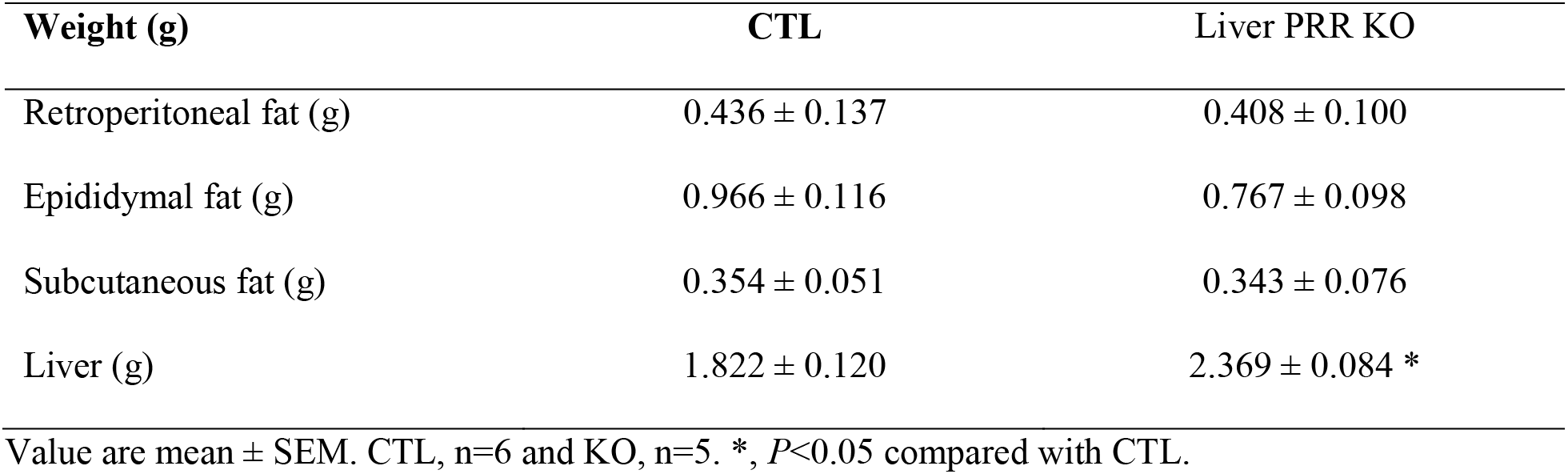
Organ weights of Liver PRR KO and control (CTL) male mice fed a standard diet

**Figure 2.**
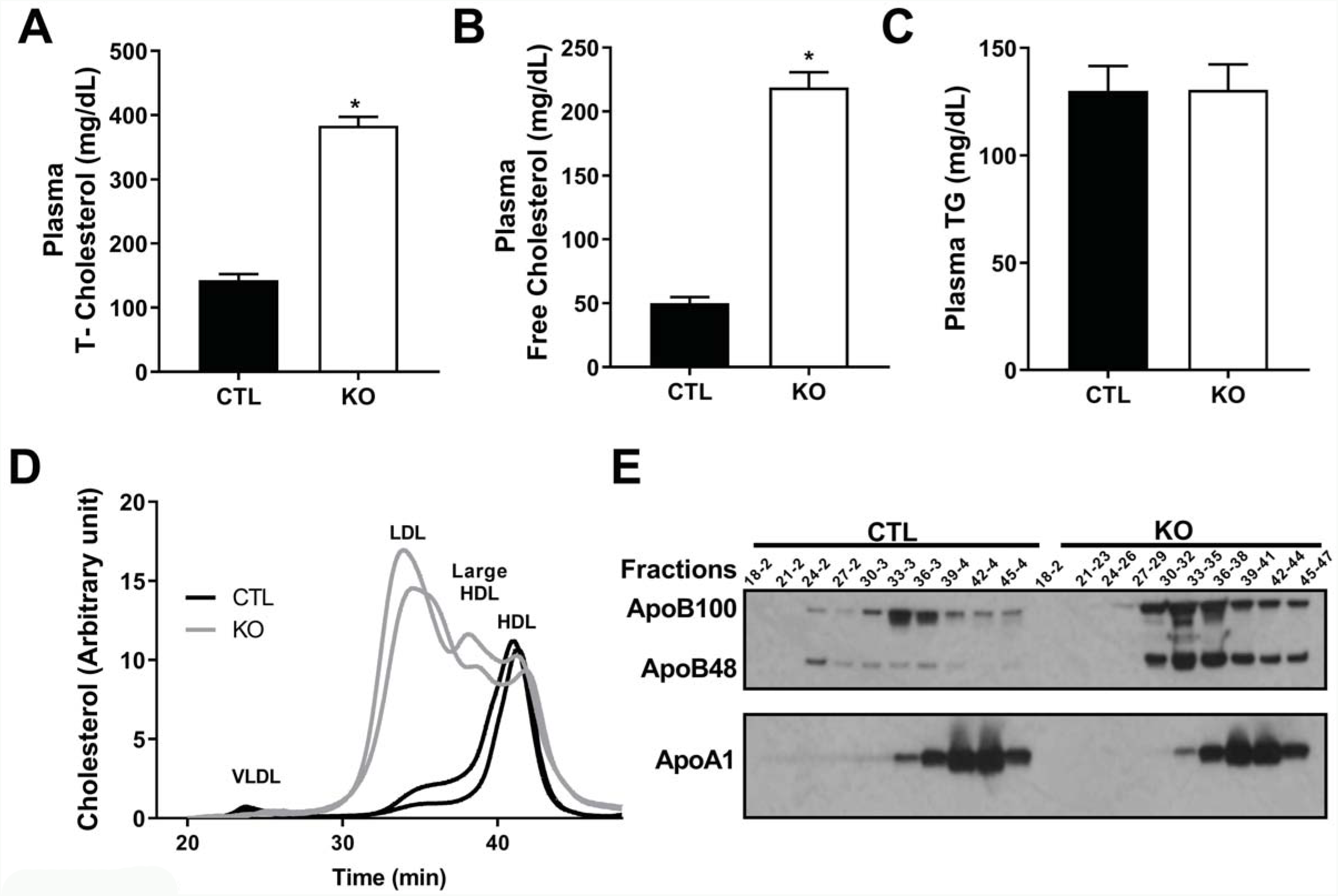
The deletion of hepatic PRR increased plasma cholesterol. (A) Plasma total cholesterol concentration in control (CTL) and liver PRR KO male mice (KO). (B) Plasma free cholesterol concentration PRR in control (CTL) and liver PRR KO male mice (KO). (C) Plasma triglycerides concentration in control (CTL) and liver PRR KO male mice (KO) (Control: n=5 and KO: n=5). Data are mean±SEM. *P<0.05 compared with vehicle. (D) Plasma cholesterol distribution determined by FPLC in two individual control (CTL) and two liver PRR KO male mice (KO). (E) Western Blot analysis of lipoprotein fractions isolated by FPLC from two pooled plasma of control (CTL) and liver PRR KO male mice (KO) (Control: n=2 and KO: n=2).

**Figure 3.**
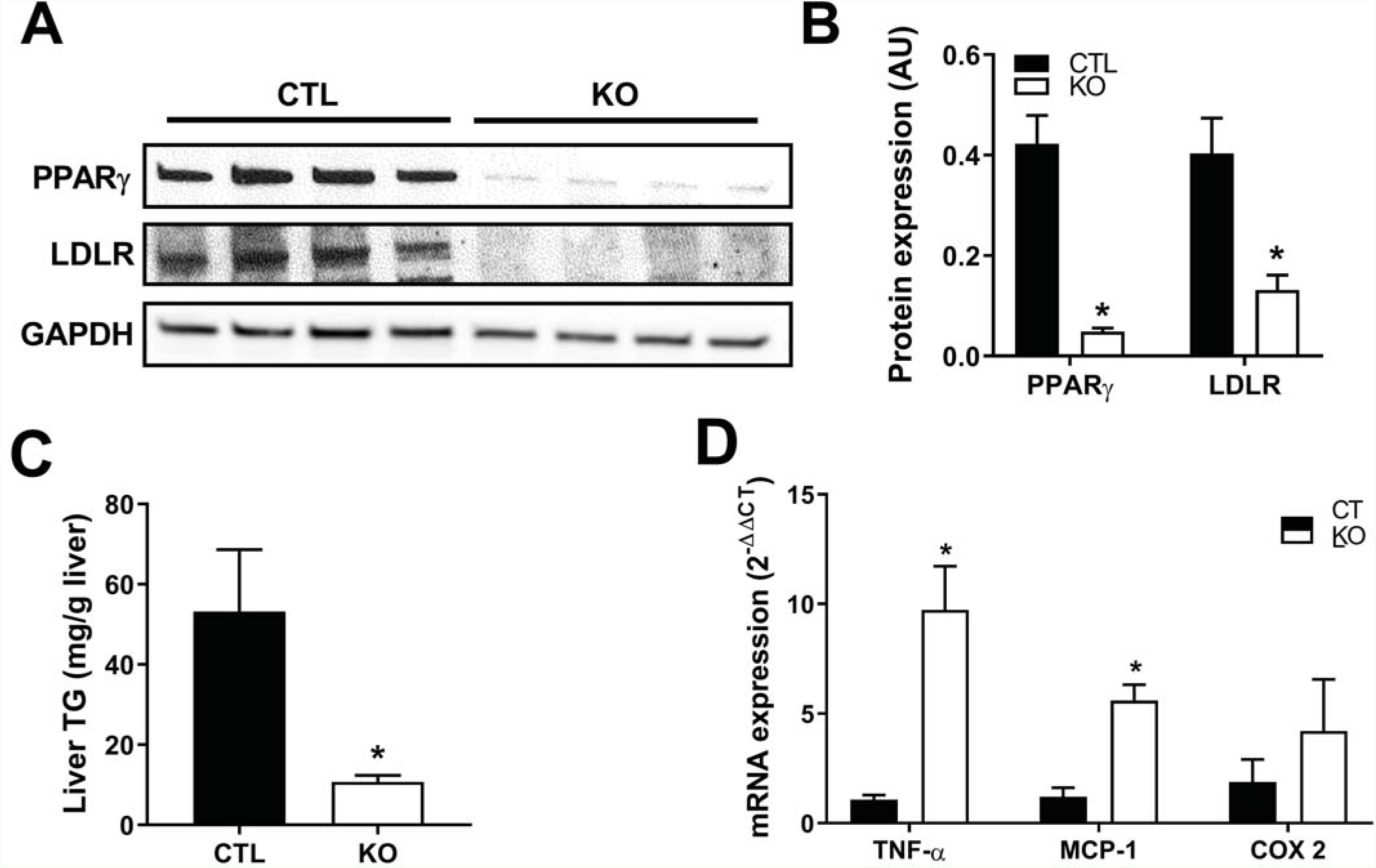
The deletion of hepatic PRR decreased hepatic triglycerides contents but promoted inflammation. (A) Representative western blot of hepatic LRLR and PPARγ proteins in control (CTL) and liver PRR KO male mice (KO). (B) Hepatic LRLR and PPARγ proteins quantification (arbitrary units) in control (CTL) and liver PRR KO male mice (KO). (C) Hepatic triglyceride contents in control (CTL) and liver PRR KO male mice (KO). (D) Hepatic mRNA expression of genes involved in inflammation in control (CTL) and liver PRR KO male mice (KO). (C: n=6 and KO: n=5). Data are mean±SEM. *P<0.05 compared with vehicle.

Because we previously showed that the deletion of PRR in adipose tissue down-regulated PPARγ and consequently prevents triglyceride accumulation in the lipid droplets of adipocytes; we aimed to determine whether hepatic PRR deletion influenced PPARγ and triglycerides contents in the liver. Indeed, PPARγ protein levels (glycosylated form, 75 KDa) and hepatic triglycerides levels decreased significantly in the liver of Liver PRR KO mice compared to CTL mice (Figure 3A, 3B and 3C).

Hepatic proinflammatory effectors TNF- α and MCP-1 mRNA levels increased significantly in Liver PRR KO mice compared to CTL mice (Figure 3D).

### The deletion of hepatic PRR elevated hepatic cholesterol likely through an up-regulation of SREBP2 and HMG CoA-reductase

The deletion of PRR in liver induced a significant increase in total cholesterol content in the liver of Liver PRR KO mice compared to CTL mice (Figure 4A, P<0.05). Furthermore, genes involved in cholesterol synthesis, especially SREBP2 and HMG CoA-reductase (HMG CoA-R), were significantly upregulated in the liver of Liver PRR KO mice compared to CTL mice (Figure 4B, P<0.05). Together our data suggested that hepatic PRR is an important regulator of endogenous cholesterol synthesis.

**Figure 4.**
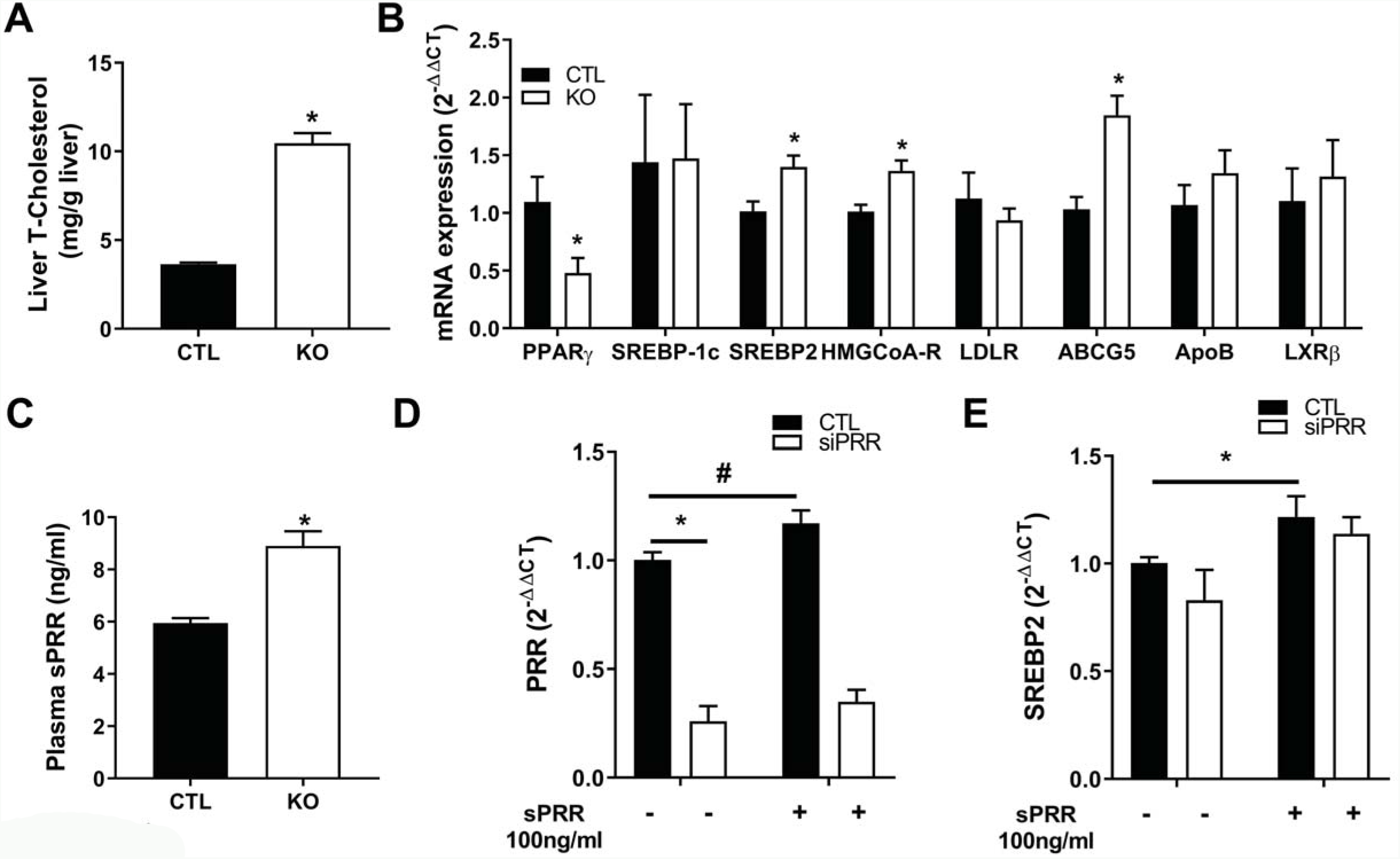
The deletion of hepatic PRR stimulated endogenous cholesterol biosynthesis pathway in the liver. (A) Hepatic cholesterol contents in control (CTL) and liver PRR KO male mice (KO) (C: n=6 and KO: n=5). (B) Expression of genes in the liver of control (CTL) and liver PRR KO male mice (KO) (C: n=6 and KO: n=5). (C) Plasma sPRR levels of PRR in control (CTL) and liver PRR KO male mice (KO). (C: n=5 and KO: n=5). The mRNA levels of PRR (D) and SREBP2 (E) in HepG2 cells transfected with siPRR and treated with vehicle or recombinant mouse sPRR-HisTag (100ng/ml) for 24h (total of 5-7 replicates from 2 experiments). Data are mean±SEM. *P<0.05 compared with vehicle.

### The deletion of hepatic PRR increased circulating sPRR, which could promote the up-regulation of hepatic SREBP2

Interestingly, Liver PRR KO mice exhibited elevated plasma sPRR levels compared to CTL mice vehicle (Figure 4C) raising the question of the relative contribution of PRR and sPRR to endogenous cholesterol synthesis. Therefore, to determine whether sPRR was involved in cholesterol synthesis, HepG2 cells were transfected with siPRR and treated with or without a mouse recombinant sPRR-His-Tag (Figure 4). PRR expression increased significantly in HepG2 cells treated with sPRR compared to HepG2 cells treated with vehicle, suggesting an autoregulatory loop in which sPRR upregulated its own gene expression (Figure 4D). The silencing of PRR in HepG2 cells did not change SREBP2 mRNA levels. However, sPRR treatment induced a significant increase of SREBP2 expression suggesting that sPRR mediated SREBP2 upregulation independently of PRR expression (Figure 4E).

### PRR and sPRR programed adipose tissue fate through up-regulation of genes involved in adipogenesis and fat mobilization

Since sPRR plasma levels doubled in Liver PRR KO mice, we next investigated the origin of this increase. Interestingly, total sPRR levels were significantly higher in the EF of Liver PRR KO mice than in CTL mice (Figure 5A) suggesting that the deletion of hepatic PRR induced a compensatory increase of plasma sPRR partially or fully originating from adipose tissue. Since we previously demonstrated that deleting PRR in adipose tissue lowers expression of genes involved in adipogenesis, lipid synthesis and trafficking (15, 19), we assessed the adipogenesis status of the adipose tissue of Liver PRR KO mice. Interestingly, the deletion of hepatic PRR led to increased PRR expression (Figure 5B) and elevated the level of PRR protein in adipose tissue (Figure 5C). In line with this increase, PPARγ, perilipin, FABP4 and ATGL gene expressions increased significantly in the EF of Liver PRR KO mice compared to CTL mice (Figure 5B). Additionally, PRR, PPARγ and perilipin protein levels were higher in the EF of Liver PRR KO mice compared to CTL mice (Figure 5C). Together our data demonstrated that the deletion of liver PRR initiated the cascade regulation of adipocyte differentiation

**Figure 5.**
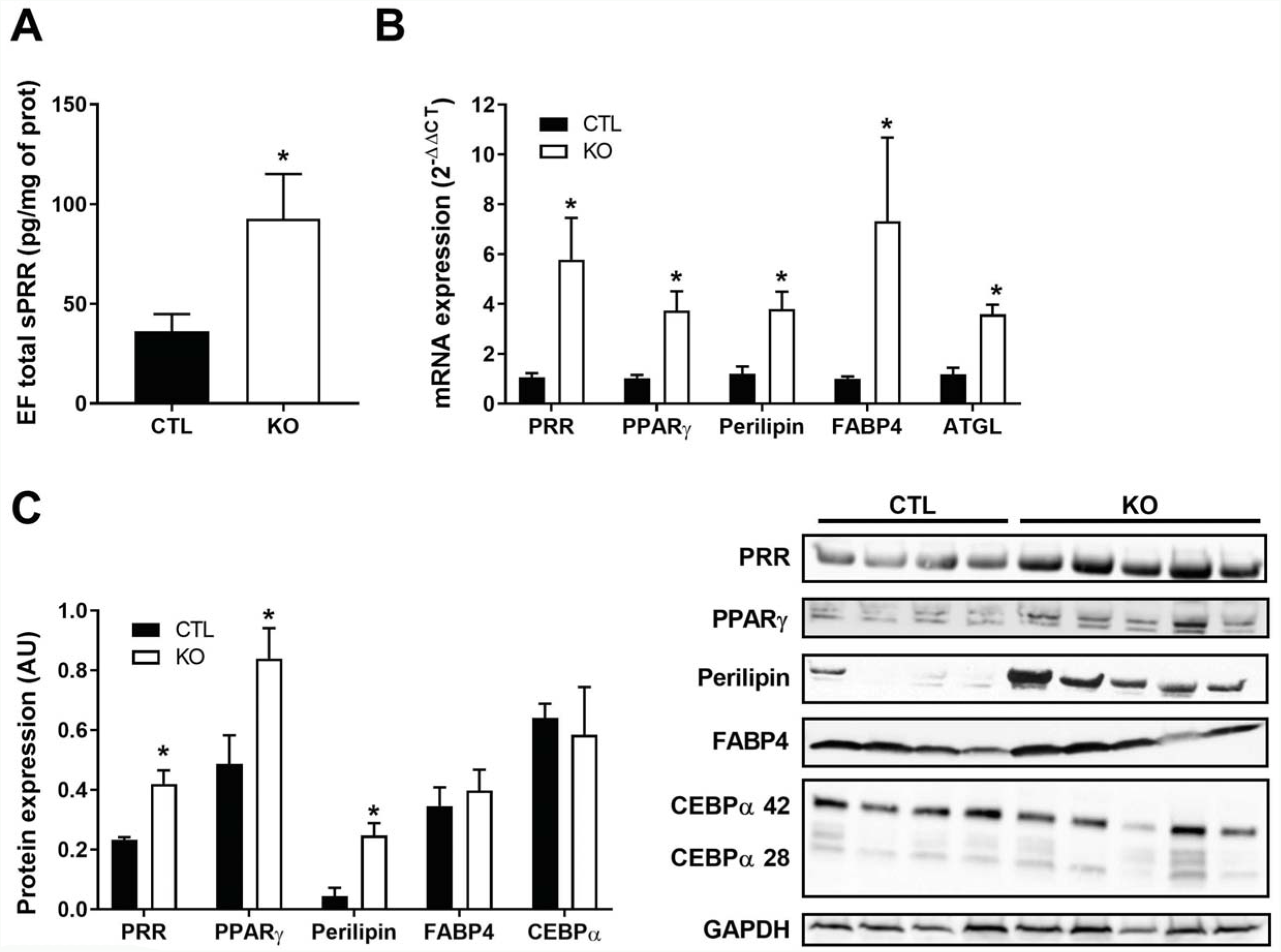
The deletion of hepatic PRR elevated total sPRR in adipose tissues and stimulated adipogenesis. (A) Total sPRR contents in the epididymal fat of control (CTL) and liver PRR KO male mice. (B) mRNA expression of genes involved in adipogenesis in epididymal fat of control (CTL) and liver PRR KO male mice. (C) Representative western blot and proteins quantification of the epididymal fat of control (CTL) and liver PRR KO male mice (KO).

To confirm the relative contribution of sPRR to the regulation of PPARγ expression, 3T3-L1 cells were treated with and without mouse recombinant sPRR-His-Tag. Our results demonstrated a dose response increase of PRR and PPARγ gene expression suggesting that sPRR might influence adipocyte differentiation by stimulating PPARγ and PRR gene expression in an autocrine manner (Figure 6A and 6B).

**Figure 6.**
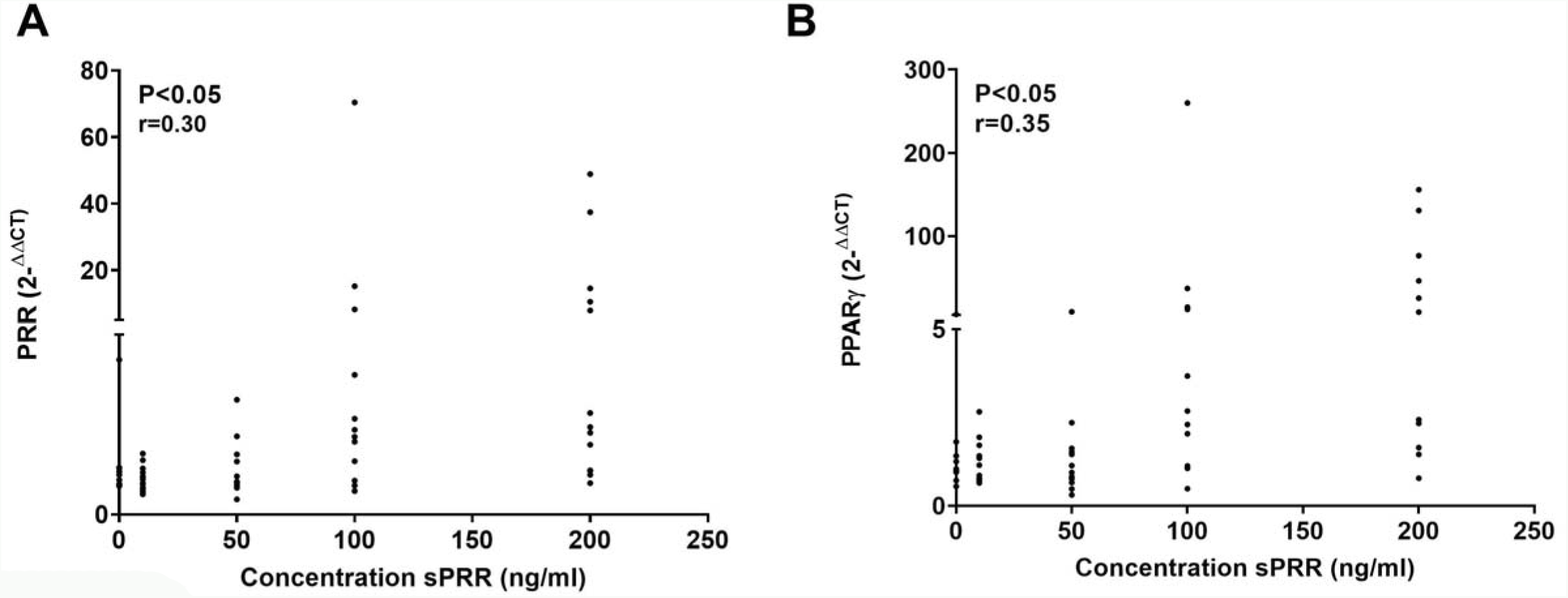
sPRR induced a dose-response increase of PRR and PPARγ. (A) The expression of PRR and (B) PPARγ genes in 3T3-L1 cells treated with vehicle or with mouse recombinant sPRR-HisTag for 24 hours. Data are mean±SEM of a total of 12 replicates from 3 experiments. r, Pearson correlation coefficient.

## Discussion

Our study demonstrated that sPRR and hepatic PRR contribute to lipid homeostasis. To our knowledge, no prior studies examined the role of sPRR in lipid homeostasis. Deleting hepatic PRR induced an increase of hepatic cholesterol, an up-regulation of SREBP2 and HMG CoA-R but also elevated circulating sPRR. Therefore, we investigated whether the stimulation of cholesterol synthesis was attributed to a lack of hepatic PRR or due to elevated levels of plasma sPRR. *In vitro* studies demonstrated that sPRR treatment up-regulated SREBP2 gene expression independently of PRR suggesting that circulating sPRR could contribute to hepatic cholesterol biosynthesis. Moreover, deleting hepatic PRR elevated total sPRR proteins levels in epididymal fat indicating that the increase of circulating sPRR likely originated from adipose tissue. Finally, we demonstrated that sPRR could participate in adipocyte differentiation through a PRR/PPARγ dependent mechanism.

The present study is in agreement with previous reports showing an association between PRR and lipoprotein metabolism (16, 17). Indeed, the silencing of PRR in HEK293, A431 and HepG2 cells impaired LDL uptake by decreasing SORT1 protein abundance, a regulator of lipid metabolism, and by reducing LDLR abundance (16). Moreover, the inhibition of hepatic PRR with a N-acetylgalactosamine (P)RR antisense oligonucleotide (ASO-PRR) reduced hepatic LDLR protein levels inducing an elevation of plasma cholesterol in mice fed a normal diet (17). Additionally, the deletion of hepatic PRR resulted in a cholesterol enrichment of LDL particles (17). In contrast, we found that hepatic cholesterol levels increased in Liver PRR KO mice whereas hepatic cholesterol levels were unchanged in ASO-PRR mice fed a normal diet. The discrepancy between the results could be attributed to the difference between the two mouse models used, ASO-PRR mice and Liver PRR KO. It is possible that the use of antisense oligonucleotides could have affected nontargeted RNAs or influenced the expression of PRR in other tissues of ASO-PRR mice (22, 23). We previously showed that the deletion of adipocyte PRR reduced adipose tissue mass through a downregulation of PPARγ gene expression and reduced genes involved in lipid transport and synthesis (15, 19). In ASO-PRR mice, PRR protein tended to decrease in the adipose tissue and the fat mass was reduced in high fat-fed ASO-PRR mice suggesting that adipogenesis might have been compromised (17). Additionally, circulating sPRR was not quantified in ASO-PRR mice and therefore the relative contribution of sPRR to the phenotype not investigated.

We previously demonstrated that the increased plasma sPRR in SD- or HF-fed adipose-PRR KO mice originated from the liver (15, 19). In the present study, we demonstrated that the increased plasma sPRR levels likely originated from the adipose tissue in liver PRR KO mice. Together, our studies suggest that a compensatory mechanism occurs between the liver and the adipose to counteract the lack of a functional tissue PRR. Hence, a liver-adipose tissue cross talk is essential for PRR regulation. In the present study, we showed that sPRR up-regulated PRR gene expression in Hep-G2 cells. This finding is consistent with our recent report showing an increase of hepatic PRR gene expression in female mice infused with sPRR (19). Interestingly, sPRR treatment cannot rescue the expression of its own gene in siPRR treated cells, suggesting that the positive feedback loop is PRR-dependent.

SREBP-2 is an essential transcription factor for de novo cholesterol synthesis, notably through its positive regulation of HMG-CoA reductase gene expression (24–26). In our study, elevated hepatic cholesterol is associated with an upregulation of SREBP-2 and HMG-CoA reductase gene expression suggesting a stimulation of cholesterol synthesis. Since circulating sPRR was elevated in Liver PRR KO, we investigated the relative contribution of PRR and sPRR to SREBP-2 regulation. sPRR treatment elevated SREBP-2 gene expression while PRR silencing had no effect on SREBP-2 transcripts, suggesting that sPRR participated in SREBP-2 regulation independently of PRR. Therefore, sPRR might be a new contributor to hepatic lipid metabolism potentially by initiating endogenous cholesterol synthesis via SREBP-2.

PRR is expressed abundantly in adipose tissue and accumulating evidences demonstrates that PRR is a master regulator of adipogenesis. In addition, the development of obesity induces an increase of PRR gene expression in adipose tissue of mice, rats, and human (15, 27, 28). Treatment with the handle region peptide, a PRR blockade, reduced fat mass and adipocyte size, as well as leptin and inflammatory cytokines levels (29). Furthermore, the silencing of PRR in 3T3-L1 cells reduced PPARγ and FABP4 gene expression indicating that PRR is an important regulator of PPARγ (15). In line with previous works, the present study indicated that the upregulation of PRR in adipose tissue was associated with an upregulation of PPARγ, FABP4 and perilipin. Additionally, sPRR promotes PRR and PPARγ gene expression in 3T3-L1 cells in a dose dependent manner. Thus, sPRR could stimulate adipogenesis through a PRR-induced PPARγ dependent mechanism. However, this stimulation does not seem enough to change the fat mass in mice fed a standard diet and a second hit (such as high fat diet) might be required to trigger this effect.

Similar to the study of Ren et al., (17), hepatic triglyceride levels and PPARγ expression were reduced in liver PRR KO mice. Previous report demonstrated that PPARγ knockdown in hepatocyte was associated with a decrease in hepatic triglycerides (30). Additionally, in AZIP Liver PPARγ KO mice, the reduction of liver triglycerides was attributed to an increase in triglycerides clearance (31). Since we showed that plasma triglycerides was not changed in liver PRR KO mice and since Ren et.al., showed a reduced hepatic triglycerides in high-fat fed ASO PRR KO, one could speculate that down-regulation of hepatic PPARγ influenced not only liver triglyceride clearance but also liver triglyceride synthesis (32).

In conclusion, we demonstrated that PRR and sPRR contributed to lipid homeostasis by stimulating regulatory pathways involved in LDLR clearance, cholesterol synthesis and adipocyte differentiation.

## Abbreviations

PRR: prorenin receptor
sPRR: soluble prorenin receptor

## Acknowledgments/grant support

We would like to thank Sierra Schlicht and Ailing Ji for their contribution to the FPLC analysis. We also are grateful to Dr. Lei Cai from Dr. Ryan Temel’s laboratory for her help with liver lipids extraction. This work was supported by National Institutes of Health Grants [R01-HL-142969]; the American Heart Association [13SDG17230008]; the National Institute of General Medical Sciences [P30 GM127211]; and the University of Kentucky, Center for Clinical and Translational Sciences [UL1TR001998].

